# The lateral hypothalamus and orexinergic transmission in the paraventricular thalamus promote the attribution of incentive salience to reward-associated cues

**DOI:** 10.1101/2020.03.01.972174

**Authors:** Joshua L Haight, Paolo Campus, Cristina E Maria-Rios, Allison M Johnson, Marin S Klumpner, Brittany N Kuhn, Ignacio R Covelo, Jonathan D Morrow, Shelly B Flagel

**Author notes:** denotes co-first author (Joshua L Haight, Paolo Campus). Corresponding author: Shelly B. Flagel, Ph.D. Associate Professor, Michigan Neuroscience Institute, Department of Psychiatry, University of Michigan. **Competing interests**: The authors report no biomedical financial interests or potential conflicts of interest.

## Abstract

**Rationale:** Prior research suggests that inputs from the lateral hypothalamic area (LHA) to the paraventricular nucleus of the thalamus (PVT) contribute to the attribution of incentive salience to Pavlovian-conditioned reward cues. However, a causal role for the LHA in this phenomenon has not been demonstrated. In addition, it is unknown which hypothalamic neurotransmitter or peptide system(s) are involved in mediating incentive salience attribution.

**Objectives:** To examine: 1) the role of the LHA in the propensity to attribute incentive salience to reward cues, and 2) the role of orexinergic signaling in the PVT on the expression of Pavlovian conditioned approach (PavCA) behavior, a reflection of incentive salience attribution.

**Methods:** Male Sprague-Dawley rats received bilateral excitotoxic lesions of the LHA prior to the acquisition of Pavlovian conditioned approach (PavCA) behavior. A separate cohort of male rats acquired PavCA behavior and were characterized as sign-trackers (STs) or goal-trackers (GTs) based on their conditioned response. The orexin 1 receptor (OX1r) antagonist SB-334867, or the orexin 2 receptor (OX2r) antagonist TCS-OX2-29, were then administered directly into the PVT to assess the effects of these pharmacological agents on the expression of PavCA behavior and on the conditioned reinforcing properties of the Pavlovian reward cue.

**Results:** Lesions of the LHA before training attenuated the development of lever-directed (sign-tracking) behaviors in the PavCA paradigm, without affecting magazine-directed (goal-tracking) behaviors. In STs, administration of the OX1r antagonist into the PVT reduced lever-directed behaviors and increased magazine-directed behaviors; while administration of the OX2r antagonist only reduced lever-directed behaviors. Further, OX2r, but not OX1r, antagonism was able to reduce the incentive motivational value of the conditioned stimulus on a conditioned reinforcement test in STs. The behavior of GTs was unaffected by orexinergic antagonism in the PVT.

**Conclusions:** The LHA is necessary for the attribution of incentive salience to reward cues and, thereby, the development of a sign-tracking conditioned response. Furthermore, blockade of orexin signaling in the PVT attenuates the incentive value of a Pavlovian reward cue. These data suggest that hypothalamic orexin inputs to the PVT are a key component of the circuitry that encodes the incentive motivational value of reward cues and promotes maladaptive cue-driven behaviors.

## Introduction

Forming stimulus-reward associations provides individuals with the fundamental capacity to identify stimuli in the environment that predict the availability of valuable resources. Such stimulus-reward associations result from Pavlovian conditioning, during which a previously neutral stimulus becomes a conditioned stimulus (CS) following repeated pairings with an unconditioned stimulus (US), such as food. In addition to acquiring predictive value, Pavlovian-conditioned stimuli (CSs) can also acquire incentive motivational value, thus becoming incentive stimuli (Bindra, 1978; Robinson & Berridge, 1993). While predictive stimuli indicate the future availability of the US, incentive stimuli evoke emotional and motivational states that render the CSs themselves ‘wanted’ (Berridge, 2001; Robinson & Flagel, 2009). Several psychiatric disorders, including substance use disorder, have been associated with an excessive attribution of incentive value to reward-cues (Berridge & Robinson, 2003; Mahler & de Wit, 2010; Versace et al., 2016; Cofresi et al., 2019; Frank et al., 2019; Hellberg et al., 2019).

Individuals differ in the extent to which they attribute incentive salience to reward-cues (Hearst & Jenkins, 1974). Exploiting this individual variability in an animal model has allowed us to parse the predictive vs. incentive qualities of reward-cues (Flagel et al., 2007; Flagel et al., 2009). When a lever-cue is repeatedly paired with a food reward, some rats, referred to as sign-trackers (STs), attribute both predictive and incentive motivational value (i.e. incentive salience) to the reward-cue, and approach and interact with the cue itself upon its presentation (Flagel et al., 2009). In addition, sign-trackers will perform a novel instrumental action for presentation of the reward-cue, even in the absence of the reward with which it was initially paired (Robinson & Flagel, 2009; Hughson et al., 2019). In contrast, other rats, referred to as goal-trackers (GTs), treat the reward-cue primarily as a predictive stimulus and approach the location of impending reward delivery upon presentation of the cue (Flagel et al., 2007; Flagel et al., 2009). Thus, while both STs and GTs attribute predictive value to the reward-cue, only STs attribute it with incentive salience (Robinson & Flagel, 2009).

The ST-GT model has provided a unique platform to investigate the neurobiology that contributes to the attribution of predictive vs. incentive value to reward-cues. Research thus far suggests that GT rats rely primarily on “top-down” cortical mechanisms to guide their behavior (Flagel et al., 2010; Paolone et al., 2013; Sarter & Phillips, 2018; Campus et al., 2019), while ST rats rely primarily on “bottom-up” subcortical circuitry (Flagel et al., 2011; Haight et al., 2017; Kuhn, 2018; Sarter & Phillips, 2018). Among the subcortical brain structures that have been shown to play a role, the paraventricular nucleus of the thalamus (PVT) has emerged as a critical mediator in regulating incentive salience attribution in STs (Flagel et al., 2011; Haight & Flagel, 2014; Haight et al., 2015; Yager et al., 2015; Haight et al., 2017; Kuhn et al., 2018; Campus et al., 2019). The PVT is a midline thalamic nucleus that is connected with cortical, limbic and motor circuitries (Chen & Su, 1990; Canteras et al., 1995; Van der Werf et al., 2002; Vertes, 2004; Kirouac et al., 2005, 2006; Vogt et al., 2008; Hsu & Price, 2009; Li & Kirouac, 2012; Li et al., 2014; Lee et al., 2015; Berendse & Groenewegen, 1990; Su & Bentivoglio, 1990; Pinto et al., 2003; Parsons et al., 2006; Parsons et al., 2007; Li & Kirouac, 2008; Vertes & Hoover, 2008).

Another subcortical brain region that seems to play a role in sign-tracking behavior is the lateral hypothalamic area (LHA) (Haight et al., 2017), which, for these purposes, refers to the lateral hypothalamus and the adjacent perifornical area. We previously demonstrated that presentation of an incentive CS evokes greater neural activity in the LHA of STs, compared to GTs, and specifically in cells that project from the LHA to the PVT (Haight et al., 2017). These data support the hypothesis that the LHA-PVT circuit plays a role in the attribution of incentive salience to reward-associated stimuli (Haight & Flagel, 2014), but the molecular identity of the cells or transmitter systems involved within this circuit remain unknown. While the inputs from the LHA to the PVT are heterogeneous, one potential candidate for mediating the incentive motivational value of reward-cues is the orexin/hypocretin system (Kelley et al., 2005; Haight & Flagel, 2014).

The orexin/hypocretin system consists of two neuropeptides, orexin-A and orexin-B, that bind to two distinct G-protein coupled receptors, orexin receptor 1 (OX1r) and orexin receptor 2 (OX2r). Orexin positive neurons originate exclusively in the LHA (de Lecea et al., 1998; Sakurai et al., 1998), and project diffusely to multiple cortical and subcortical brain regions (Marcus & Elmquist, 2006). While known primarily for its role in arousal, sleep and feeding behavior, orexinergic transmission has long been implicated in cue-motivated and addiction-related behaviors (Mahler et al., 2012; Petrovich et al., 2012; Cason & Aston-Jones, 2013; Sakurai, 2014; Cole et al., 2015; Keefer et al., 2016). Recent evidence suggests that the role of orexin in mediating such behaviors is, in part, localized to the PVT (Li et al., 2011; Martin-Fardon & Boutrel, 2012; James & Dayas, 2013; Matzeu et al., 2014; Barson et al., 2015; Matzeu et al., 2016; Matzeu & Martin-Fardon, 2018), which receives dense orexinergic projections (Kirouac et al., 2005; Lee et al., 2015).

To better examine the role of the LHA and orexinergic signaling in the PVT on incentive motivational processes, we performed two experiments. In Experiment 1, we performed bilateral excitotoxic lesions of the LHA in male Sprague-Dawley rats before training them in a Pavlovian conditioning paradigm. In Experiment 2, after rats had acquired Pavlovian conditioned approach behavior, we administered either the OX1r antagonist, SB-334867 (Experiment 2a), or the OX2r antagonist, TCS-OX2-29 (Experiment 2b), into the PVT and assessed the effects on the expression of sign- and goal-tracking behavior, and on the conditioned reinforcing properties of the Pavlovian-conditioned food cue. We hypothesized that lesions of the LHA, and blockade of orexin signaling in the PVT, would attenuate the attribution of incentive salience to a Pavlovian conditioned food cue and thereby the expression of sign-tracking behavior.

## Materials and methods

All procedures were approved by the University of Michigan Institutional Animal Care and Use Committee, and all experiments were conducted in accordance with the National Academy of Sciences Guide for the Care and Use of Laboratory Animals: Eighth Edition, revised in 2011.

### Housing

Male Sprague-Dawley rats (Charles River Saint-Constant, Canada and Raleigh, NC, USA) were used. Rats were housed in a climate-controlled room (22±2 °C) with a 12-hour dark-light cycle (lights on at 06:00 or 07:00 depending on daylight savings time). All rats had ad-libitum access to food and water for the duration of the experiments. Behavioral testing took place during the light cycle between 11:00 and 17:00.

### Surgeries

Surgeries were performed under aseptic conditions. A surgical plane of anesthesia was induced with inhalation of 5% isoflurane, and anesthesia was maintained throughout the procedure with inhalation of 1-2% isoflurane. Prior to surgeries, while under anesthesia, rats received an injection of carprofen (5mg/kg, s.c.) for analgesia and were further prepared for surgeries by shaving the scalp and applying betadine (Purdue Products, Stamford, CT) followed by 70% alcohol as an antiseptic. Rats were then placed into a stereotaxic frame (David Kopf instruments, Tujunga, CA or Stoelting, Wood Dale, IL) and a small incision was made on the scalp to expose the skull. The skull was leveled within +/− 0.1 mm using bregma and lambda coordinates, and small holes were drilled above the regions of interest, as described below.

### Pavlovian conditioned approach (PavCA) apparatus

PavCA training occurred inside Med Associates chambers (St. Albans, VT, USA; 30.5□×□24.1□×□21□cm) located in sound-attenuating cabinets with a ventilation fan to create background noise. Each chamber contained a food magazine located in the center of one wall approximately 3 cm above the grid floor, connected to an automatic pellet dispenser. Each time the pellet dispenser was triggered, one 45-mg banana-flavored dustless pellet (Bio-Serve, Flemington, NJ) was delivered into the food cup. A retractable backlit metal lever was located either to the left or right of the food magazine, approximately 6 cm above the grid floor. A house light was located on the wall opposite to the food magazine and lever, approximately 1 cm from the top of the chamber. Magazine entries were recorded upon break of a photo-beam located inside the magazine and lever contacts were registered upon deflection of the lever, which required a minimum of 10 g of force. Behavioral data were collected using Med Associates’ Med PC software.

### PavCA Procedure

PavCA procedures were the same as described previously (Campus et al., 2019; Hughson et al., 2019). For two days prior to behavioral training rats were briefly handled by the experimenters in the housing room, and a small scoop (~25) of banana-flavored pellets were delivered in the home cage, to familiarize rats to the experimenters and to the novel food. Following these two days, all rats underwent one pretraining session, followed by 7 PavCA sessions. The pretraining session consisted of 25 trials in which a food pellet was delivered into the food magazine on a variable time (VT) 30-s schedule (range 0-60 s). Prior to the start of the session, each food magazine was baited with 3 banana-flavored pellets, to direct the rats’ attention to the location of reward delivery. The lever remained retracted for the entirety of the session, which lasted an average of 12.5 minutes. Rats typically consumed all of the pellets delivered into the food cup during the pretraining session. After pre-training, rats underwent one daily session of PavCA training for 7 consecutive days. Each PavCA training session consisted of 25 trials under a VT-90 s schedule (range 30-150 s). During each trial, an illuminated lever was inserted into the chamber for 8 s, and was followed by the delivery of a banana-flavored food pellet into the food magazine upon lever retraction. The start of PavCA training was signaled by illumination of the house light and lasted an average of 40 minutes. For each PavCA session, the number of lever contacts and head entries into the food magazine, the probability of contacting the lever or entering the food magazine, and the latency to contact the lever or to enter the food magazine during each trial were recorded or calculated.

## Detailed Methods

### Experiment 1-Effects of LHA lesion on the acquisition of Pavlovian conditioned approach behavior

#### Subjects

Sixteen adult male Sprague-Dawley rats (Charles River Saint-Constant, Canada), weighing an average of 400 g (~11-12 weeks of age) at the time of experimentation were used. Rats were pair-housed upon arrival. Following three days of acclimation to the colony room, the rats went through a three-day fear-conditioning pilot experiment (data not included) and were then allowed to rest in the colony room for two weeks. After the two-week rest period, lesion and sham surgeries were performed. Following surgery, rats were single-housed for the remainder of the experiment, and allowed to rest for an additional two weeks prior to the start of PavCA training. The fear-conditioning pilot experiment was unlikely to affect the results of the current experiment, due to the substantial rest periods (4 weeks total) and the fact that all subjects were counter-balanced across treatment groups.

#### Drugs

Excitotoxic lesions were performed using N-methyl-D-aspartate (NMDA; #M3262; Sigma-Aldrich, Inc.; St. Louis, MO). NMDA was dissolved in sterile saline and injected bilaterally in the LHA at a 90 mM concentration (pH□=□7.34-7.36).

#### Lesion surgery and PaVCA training

Excitotoxic lesions (Figure 1a) were performed by lowering one-barrel stainless steel guide-cannulas (26-gauge; Plastics One, Inc.; Roanoke, VA) bilaterally into the LHA at the following coordinates relative to bregma: AP: −2.2 mm, ML: +/−1.7 mm, DV: −8.1 mm. A stainless steel injector that projected 1mm beyond the guide cannula (33-gauge; Plastics One, Inc.; Roanoke, VA) was inserted into each cannula and connected with PE-20 tubing to a microsyringe (5 μL; Hamilton Company; Reno, NV) mounted in an infusion pump (Harvard Instruments; Holliston, MA). NMDA injections occurred over the course of 4 min at a rate of 0.15 □l/min, for a total delivered volume of 0.6 □l. The injector was left in place for two minutes following injection to allow for diffusion. Sham surgeries were done by performing the same incisions and by drilling holes over the LHA, but cannulas were not inserted and no injections were delivered. Following surgery, the incisions were sealed using surgical clips.

**Figure 1.**
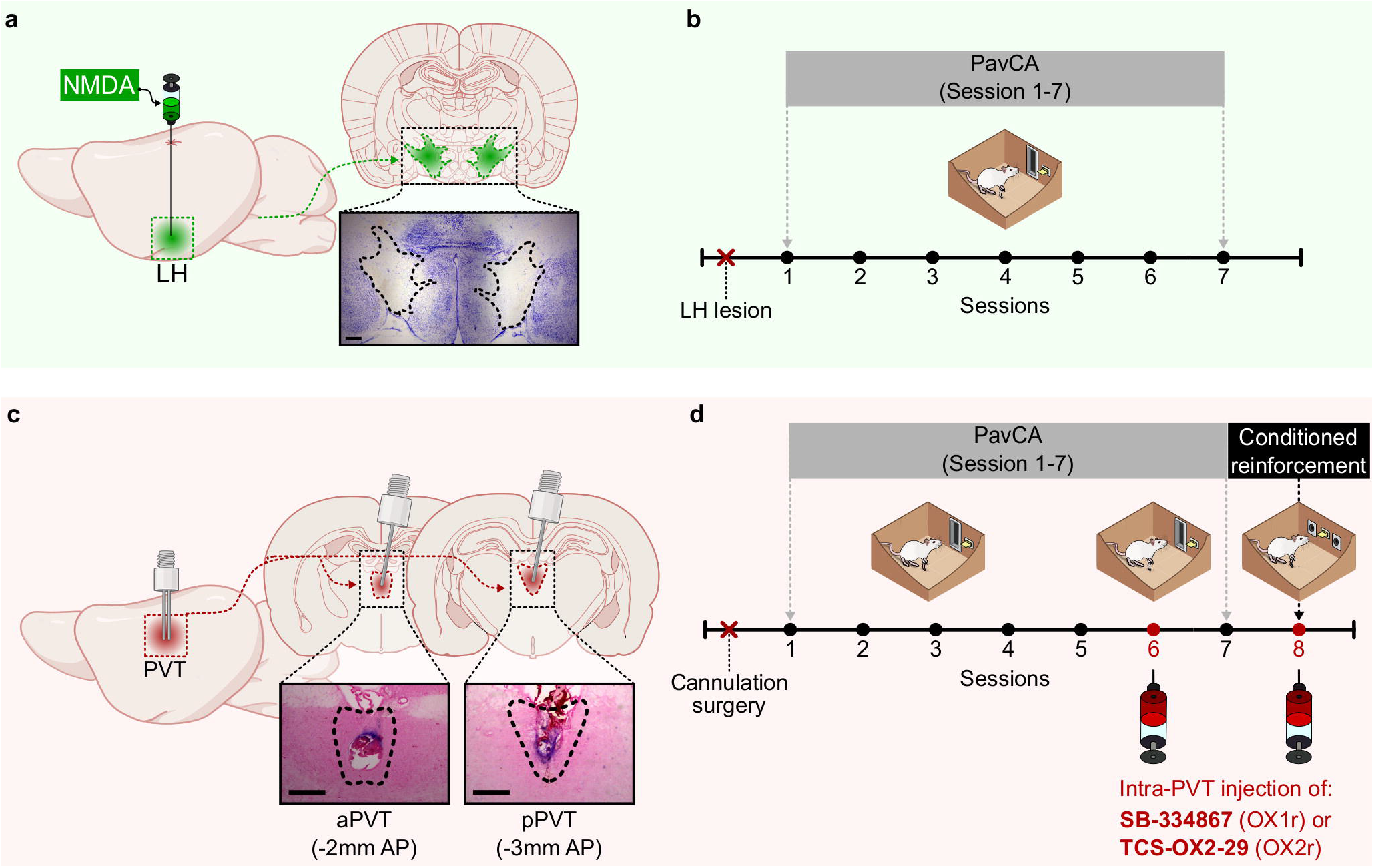
Experimental timelines. Experiment 1: **a)** Schematic of the excitotoxic lesion of the LH and photomicrograph of a representative lesion, and **b)** experimental timeline. Rats received 0.6 □l of NMDA in the right and left LHA. Sham rats underwent the same surgery but no injection was performed. After recovery from surgery, rats were trained in a Pavlovian Conditioned Approach (PavCA) paradigm for 7 consecutive sessions. Experiment 2: **c)** Schematic of the cannulation surgery to target the aPVT and pPVT and representative images of aPVT and pPVT cannulation sites, and **d)** experimental timeline. After recovery from surgery, rats were trained in a Pavlovian Conditioned Approach (PavCA) paradigm for 5 consecutive sessions (Acquisition) and phenotyped as sign-(STs) or goal-trackers (GTs). On the following day rats went through a PavCA test session (session 6) in order to assess the effects of OX1 or OX2 receptor antagonism in the PVT on the expression of Pavlovian conditioned approach behavior. The next day, all rats went through an additional PavCA session (session 7) with no drug or vehicle infusions to assess any lasting effects of drug infusion on PavCA behavior. Following the completion of PavCA training, all subjects were tested in a conditioned reinforcement (CRT) paradigm, in order to assess the effects of OX1 or OX2 receptor antagonism in the PVT on the conditioned reinforcing properties of the lever-CS.

#### Tissue processing

Rats were deeply anesthetized with 5% inhaled isoflurane. Following anesthetization, brains were extracted and flash frozen in isopentane cooled with dry ice. The frozen brains were subsequently sectioned in the coronal plane (40 □m) using a cryostat (Leica Biosystems Inc, Buffalo Grove, IL), mounted onto SuperFrost Plus microscope slides (Fisher Scientific), soaked in 4% formaldehyde for 30 minutes, and counterstained with cresyl violet. Sections containing the LHA were assessed for lesion accuracy by an experimenter blind to the experimental groups using a DM1000 light microscope coupled to an ICC50 HD camera (Leica-Microsystems, Wetzlar, GER).

#### Statistical analyses

To assess differences in the acquisition of PavCA behavior, a linear mixed effects model with a restricted maximum likelihood estimation method was used. Session was used as the repeated variable and Group (Sham vs. Lesion) as the between-subject variable. Lever-directed behaviors (lever contacts, probability to contact the lever and latency to contact the lever) and magazine-directed behaviors (magazine entries during the CS period, probability to enter the magazine during the CS period and latency to enter the magazine during the CS period) were used as dependent variables. Before choosing the final model, several covariance structures were explored for each one of the dependent variables, and the best-fitting model was chosen by selecting the lowest Akaike Information Criterion (AIC) (Verbeke & Molenberghs, 2009; Duricki et al., 2016). All statistical analyses were performed using IBM SPSS Statistics 26 (IBM, Armonk, NY, USA). Alpha was set at 0.05. When significant main effects or interactions were detected, Bonferroni post-hoc comparisons were performed. Graphic representations of the data were created with Prism 8 (Graphpad Software, San Diego, CA).

### Experiment 2-Effects of the pharmacological antagonism of orexin 1 or orexin 2 receptors on the expression of sign-tracking behavior and on the conditioned reinforcing properties of a Pavlovian reward-cue

#### Subjects

A total of 230 male Sprague Dawley rats (Charles River Saint-Constant, Canada and Raleigh, NC, USA) were used for Experiment 2. Rats were 275-325 g (7-9 weeks old) at the time of arrival and were initially pair-housed and allowed to acclimate to the housing room for at least 7 days prior to the surgery. Following cannula implantation surgery (described below), rats were single-housed for the remainder of the study to avoid damage to the implanted cannulas. As we were interested only in assessing STs (n=116) and GTs (n=62), rats with a PavCA index between −0.3 and +0.3 (intermediate rats, n = 52) were excluded from Experiment 2. Of the remaining 178 rats, 88 were excluded from the analysis due to missed cannula placement in either the aPVT or the pPVT. In addition, 9 rats did not complete the experiment due to health or technical issues, resulting in a final n of 81 rats (52 STs, 29 GTs). Of the final 81 rats, 47 (29 STs, 18 GTs) were used for the OX1r antagonism study (Experiment 2a) and 34 (23 STs, 11 GTs) were used for the OX2r antagonism study (Experiment 2b). Data for Experiment 2a were collected across 3 rounds of testing, while data for Experiment 2b were collected across 2 rounds of testing.

#### Drugs

To block orexin 1 receptors, the selective OX1r antagonist SB-334867 (SB; Lots 11B/185592, 11B/186281, Tocris Bioscience, Avonmouth, Bristol, UK) was dissolved in 100% dimethyl sulfoxide (DMSO) at a concentration of 15 μg per 300 nl. To block orexin 2 receptors, the selective OX2r antagonist TCS-OX2-29 (TCS; Lots 2A/179223, 2B191601, Tocris Bioscience) was dissolved in 0.9% sterile saline at a concentration of 15 μg per 300 nl.

#### Guide cannula implantation

For both Experiment 2a and 2ba stainless steel double cannula aligned along the anterior-posterior axis (1 mm center to center gap, cut 6 mm below pedestal, 26 gauge, part # C235G-1.0-SP, Plastics One, Roanoke, VA) was implanted with the stereotaxic arm angled at 10° towards the midline to target the aPVT and the pPVT (Figure 1b), at the following coordinates relative to bregma: AP −2.0 mm, ML −1.0 mm, DV −4.7 mm (aPVT) and AP −3.0 mm, ML −1.0 mm, DV −4.7 mm (pPVT). Four screws were then implanted in the skull, and the cannula was fixed in place using dental cement. Once the cement was dry, the incision was closed around the cement with stainless steel wound clips. In addition, the cannula was plugged with a dummy injector that was flush with the end of the cannula, and covered with a dust cap. Following cannulation surgeries rats were allowed to recover a minimum of 7 days prior to any behavioral testing.

#### PavCA training

All subjects from Experiments 2a and 2b went through 1 session of pretraining, followed by 7 sessions of PavCA training as described above (for experimental timeline, see Figure 1d). Prior to pretraining and the first 5 PavCA sessions, rats were transported to a separate room and handled by the experimenters, increasing in time from approximately 30 seconds (pretraining) to 4 minutes (PavCA session 5). In addition, during the handling prior to sessions 4 and 5, all rats had their dust caps screwed on and off, in order to acclimate them to the infusion procedure.

#### Classification of rats into STs or GTs

Following session 5 of PavCA training, rats were classified as STs or GTs based on their average PavCA Index scores (Meyer et al., 2012) from sessions 4 and 5. The PavCA Index is a composite score that is used to measure the degree to which an individual’s behavior is directed towards the lever-CS or food cup (location of US delivery) using three different metrics: response bias [(total lever contacts – total food cup contacts) / (sum of total contacts)], probability difference score [Prob(lever) – Prob(food cup)], and latency difference score [-(lever contact latency – food cup entry latency) / 8 seconds]. These three measures were averaged together and rounded to the nearest tenth of a decimal place to create the PavCA Index score, which ranges from −1.0 to 1.0, with −1.0 representing an individual whose behavior is directed solely towards the food cup (i.e. goal-tracker), and 1.0 representing an individual whose behavior is directed solely towards the lever-CS (i.e. sign-tracker).

#### Intra-PVT infusions and PavCA test

Following ST/GT classification, rats were split into experimental and control groups counterbalanced based on their PavCA Index score. On the following day rats went through a PavCA test session (session 6) in order to assess the effects of OX1r (Experiment 2a) or OX2r (Experiment 2b) antagonism in the aPVT and pPVT on the expression of PavCA behavior. Prior to the test session, each rat’s dust cap and dummy cannula was removed, and a double injector protruding 1 mm beyond the guide cannula was inserted (final infusion coordinates AP −2.0 mm, ML −1.0 mm, DV −5.7 mm and AP −3.0 mm, ML −1.0 mm, DV −5.7 mm). The injector was connected via P50 tubing to two 1 μl Hamilton syringes housed in a Harvard Apparatus double syringe pump. Rats from Experiment 2a were infused with either SB-334867 (SB) or 100% dimethyl sulfoxide (DMSO) as control vehicle. Rats from Experiment 2b were infused with either TCS-OX2-29 or 0.9% saline (SAL) as a control vehicle. Infusions occurred simultaneously in the aPVT and pPVT and lasted 2 minutes, at a flow rate of 150 nl per minute, for a total infusion volume of 300 nl per injection site.

After the end of the infusion, the injector was left in place for 2 additional minutes to allow diffusion. An experimenter gently held each rat for the 4-minute duration of the infusion/diffusion period. After the injector was removed, the dummy cannula and dust cap were replaced, and the rat was placed back into its home cage and transported to the testing room, where it sat for 10 minutes under red light prior to being placed in the chambers for the PavCA test. The next day, all rats went through an additional PavCA session (session 7) with no drug or vehicle infusions to assess any lasting effects of drug infusion on PavCA behavior.

#### Conditioned Reinforcement Test (CRT)

Following the completion of PavCA training, all subjects were tested in a conditioned reinforcement (CRT) paradigm, in order to assess the effects of intra-PVT OX1r (Experiment 2a) or OX2r (Experiment 2b) antagonism on the conditioned reinforcing properties of the lever-CS. Injection procedures before CRT were the same as described above, and all treatment groups remained consistent from the PavCA experiment to the CRT experiment. For the CRT test, the chambers were rearranged so that the food cup and pellet dispenser were removed, and the lever-CS was moved to the center of the wall. Two nose poke ports were then installed to the right and left of the lever. The nose port installed opposite the previous position of the lever-CS was designated the “active” nose port, and pokes into this port resulted in the brief 2-second presentation of the illuminated lever-CS on a fixed-ratio 1 schedule. Pokes into the other nose port, designated the “inactive”, did not result in lever-CS presentation. Once the rats were placed into the test chambers, the house light remained off for 1 minute. After the 1-minute acclimation period, the house light was illuminated, and the CRT test session began. The session lasted 40 minutes, and the Med PC software program recorded the following measures for analysis: 1) the number of pokes into the active nose port, 2) the number of pokes into the inactive nose port, and 3) the number of lever contacts.

#### Tissue processing

Rats were deeply anesthetized with an intraperitoneal injection of ketamine (90 mg/kg i.p.) and xylazine (10 mg/kg i.p.) and transcardially perfused with ~200 mL of room temperature 0.9% saline, followed by ~200 mL of room-temperature 4% formaldehyde (pH=7.3-7.4, diluted in 0.1M sodium phosphate buffer; Fisher Scientific, Hampton, NH). Brains were then extracted and post-fixed overnight in 4% formaldehyde at 4°C. Brains were cryoprotected over three nights in graduated sucrose solutions (10%, 20%, and 30%, dissolved in 0.1M sodium phosphate buffer with a pH=7.3-7.4) at 4°C. Following cryoprotection, brains were encased in Tissue-Plus O.C.T. (Fisher HealthCare, Houston, TX), frozen using dry ice and subsequently sectioned in the coronal plane (40 □m) using a cryostat (Leica Biosystems Inc, Buffalo Grove, IL). Brain slices containing the PVT were collected into well plates containing cryoprotectant and stored at −20 °C before being mounted onto SuperFrost Plus microscope slides (Fisher Scientific), and counterstained with Eosin-Y. Sections were assessed for accuracy of cannula placement using a DM1000 light microscope coupled to an ICC50 HD camera (Leica-Microsystems, Wetzlar, GER) by an experimenter blind to the experimental groups.

#### Statistical analyses

Similar to Experiment 1, linear mixed effects models with a restricted maximum likelihood estimation method were used to assess differences in the acquisition and expression of PavCA behavior. For acquisition of PavCA behavior across all subjects, Session (1-5) was used as the repeated variable and Phenotype (ST vs. GT) was used as between-subjects variable. Lever-directed behaviors (lever contacts, probability to contact the lever and latency to contact the lever) and magazine-directed behaviors (magazine entries during the CS period, probability to enter the magazine during the CS period and latency to enter the magazine during the CS period) were used as dependent variables. Similarly, PavCA index scores during session 1-5 of PavCA training were analyzed across all subjects using a linear mixed effect model. For this analysis, session (1-5) was used as the repeated variable and Phenotype (ST vs. GT) as the between-subject variable. In addition, to ensure that subjects were counterbalanced between different experimental groups, baseline differences in lever- and magazine-directed behaviors during session 5 (prior to treatment) and the average PavCA index scores from sessions 4-5 were assessed using a two-way ANOVA, with Phenotype (STs vs. GTs) and Treatment (DMSO vs. SB for Experiment 2a; SAL vs. TCS for Experiment 2b) as independent variables.

To assess the effects of OX1r or OX2r antagonism in the PVT on the expression of PavCA behavior, the effect of Treatment was assessed within each phenotype separately. Linear mixed effects models with a restricted maximum likelihood estimation method were used, with Session (5-7) as the repeated variable and Treatment (SB vs. DMSO for Experiment 2a; TCS vs. SAL for Experiment 2b) as the between-subject variable. Lever contacts and magazine entries were used as the dependent variables. For all linear mixed effects models, the covariance structure that was the best fit was chosen based on the lowest AIC.

To assess the effects of OX1r and OX2r antagonism in the PVT on the behaviors expressed during CRT, nose pokes, lever contacts and incentive value index (Campus et al., 2019; Hughson et al., 2019) were analyzed within each phenotype. Nose pokes were analyzed using a two-way ANOVA with nose port (Active vs. Inactive) and Treatment (DMSO vs. SB for Experiment 2a; SAL vs. TCS for Experiment 2b) as independent variables. Differences in lever contacts and in the incentive value index were analyzed using an unpaired t-test. The incentive value index was calculated as previously described (Hughson et al., 2019) by using the following formula: [(Active Nosepokes + Lever Contacts) – Inactive Nosepokes]. All statistical analyses were performed using IBM SPSS Statistics 26 (IBM, Armonk, NY, USA). Alpha was set at 0.05. When significant main effects or interactions were detected, Bonferroni post-hoc comparisons were performed. Graphic representations of the data were created with Prism 8 (Graphpad Software, San Diego, CA).

## Results

### Experiment 1-Effects of LHA lesion on the acquisition of Pavlovian conditioned approach behavior

#### Lesion verification

Sections containing the LHA were screened for accuracy of lesion placement. All subjects in the lesion group showed evidence of excitotoxic damage to the LHA, with lesions generally spanning from −2.12 to −2.80 AP, relative to bregma. A representative image of an LHA lesion can be found in Figure 1a, and the spread of LHA lesions for all experimental rats can be found in Supplemental Figure 1.

#### Lesions of the LHA impair the acquisition of sign-tracking but have no effect on goal-tracking

To assess the effects of a LHA lesion on the acquisition of sign-and goal-tracking behavior, lever-directed behaviors (lever contacts, probability to contact the lever and latency to contact the lever) and magazine-directed behaviors (magazine entries during the CS period, probability to enter the magazine during the CS period and latency to enter the magazine during the CS period) were analyzed across 7 sessions of PavCA training. Relative to lesioned rats, control (sham) rats exhibited a greater number of lever contacts (*F*_6,22.118_ = 2.565, *p* = 0.049, Figure 2a), a greater probability to contact the lever (*F*_6,36.592_ = 3.000, *p* = 0.017, Figure 2b), and had a lower latency to approach the lever (*F*_6,30.768_ = 3.640, *p* = 0.008, Figure 2c). Post-hoc analyses revealed a significant difference between sham and lesioned rats from Session 3-7 (p < 0.044) for all lever-directed behaviors. There was not a significant effect of group for magazine entries (*F*_6,23,484_ = 0.861, *p* = 0.537, Figure 2d), probability to enter the magazine (*F*_6,78.090_ = 0.501, *p* = 0.806, Figure 2e) or for latency to enter the magazine (*F*_6,18.702_ = 0.411, *p* = 0.862, Figure 2f). Thus, lesions of the LHA impair the acquisition of lever-directed behaviors, without affecting goal-directed behavior. These data highlight a role for the LHA in the attribution of incentive salience to a reward-cue.

**Figure 2.**
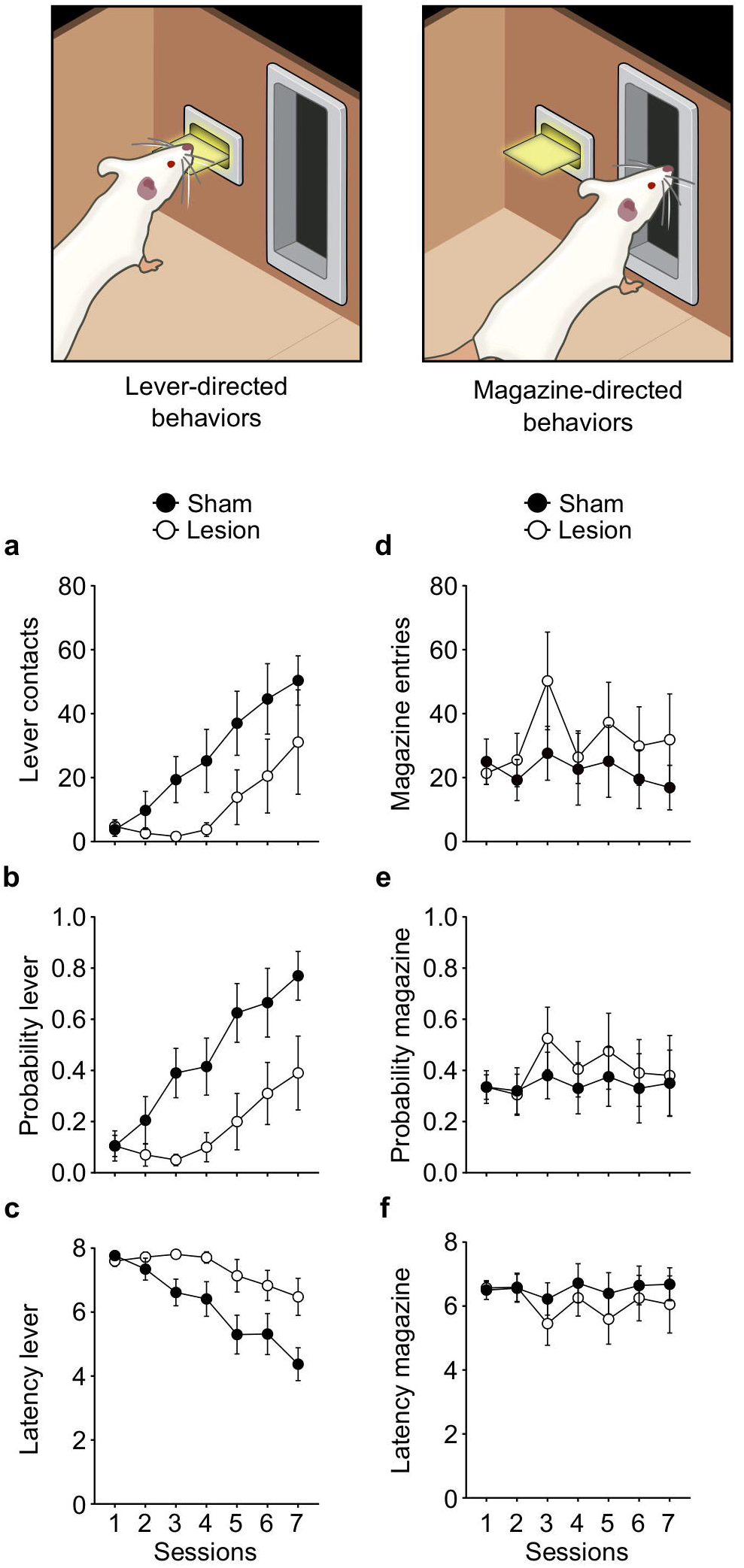
Effects of LHA lesions on the acquisition of lever-directed (sign-tracking) and magazine-directed (goal-tracking) behaviors in Experiment 1. Mean ± SEM for **a**) Number of lever contacts, **b)** probability to contact the lever, **c)** latency to contact the lever, **d)** number of magazine entries, **e)** probability to enter the magazine and **f)** latency to enter the magazine across 7 sessions. There was a significant effect of treatment (Sham vs. lesion) on sign-tracking (left column; P<0.05), but not goal-tracking behaviors (right column). N = 8/group.

### Experiment 2-Effects of the pharmacological antagonism of orexin 1 or orexin 2 receptors on the expression of Pavlovian conditioned approach behavior and on the conditioned reinforcing properties of a Pavlovian reward-cue

#### Cannula placement verification

Cannula placements were verified for all subjects to ensure accuracy of drug infusions into the anterior and posterior PVT. Subjects with injector tracts abutting the dorsal border of the PVT, within the PVT, or immediately adjacent to the lateral or ventral borders of the PVT were considered accurate injections, and remained in the study (for an example of acceptable cannula placement, see Figure 1b). Subjects with at least one injection that did not touch the top of the PVT, or that was completely contained within the medio dorsal nucleus or the habenula, were excluded from the study, so that only subjects with both successful anterior and posterior PVT infusions were included. As stated above, 88 animals were excluded from Experiment 2, due to missed cannula placements in either the anterior PVT, posterior PVT, or both, and are not included in the analyses below.

#### Acquisition of PavCA behaviors

Similar to previous reports (Robinson & Flagel, 2009; Meyer et al., 2012), there was variation in the conditioned responses acquired following 5 sessions of PavCA. Rats that directed their behavior towards the food magazine were classified as GTs, with average PavCA index scores from sessions 4 and 5 ranging between −0.3 and −1.0 (Figure 3g). Rats that displayed lever-directed behavior were classified as STs, with PavCA Index scores ranging from 0.3 to 1.0 (Figure 3g). Animals with a PavCA index scores ranging from −0.3 to 0.3 (intermediate rats), were not included in the study

**Figure 3.**
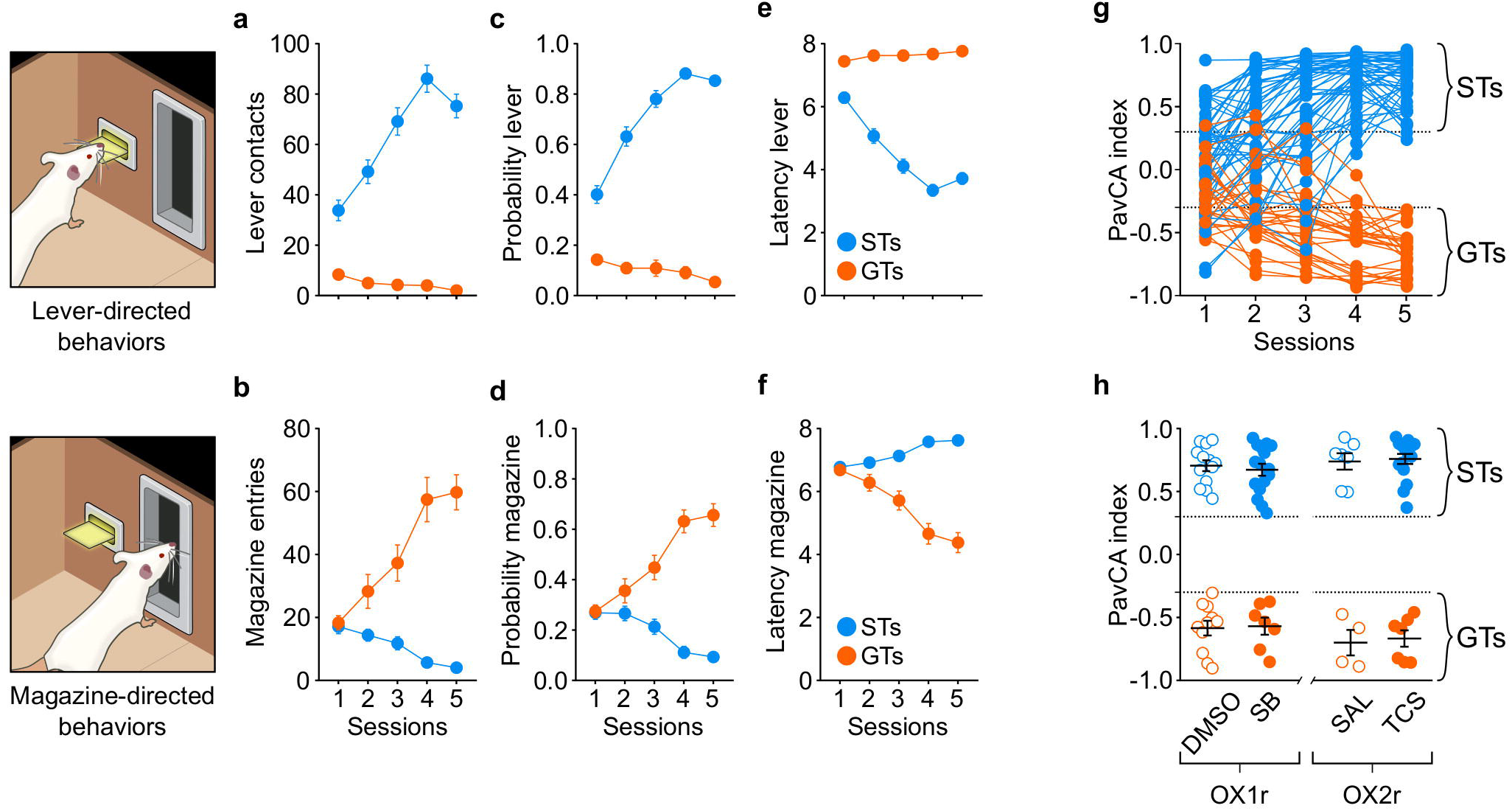
Acquisition of lever-directed (sign-tracking) and magazine-directed behaviors (goal-tracking) for Experiment 2. **a-f)** Mean ± SEM for **a**) Number of lever contacts, **b)** number of magazine entries, **c)** probability to contact the lever, **d)** probability to enter the magazine, **e)** latency to contact the lever and **f)** latency to enter the magazine. **g)** Individual Pavlovian conditioned approach (PavCA) index scores during the 5 sessions of Pavlovian conditioning. PavCA scores from session 4 and 5 were averaged to determine the behavioral phenotype. Rats with a PavCA score <−0.3 were classified as goal-trackers (GTs), rats with a PavCA score >+0.3 were classified as sign-trackers (STs; n= 52 STs, 29 GTs). **h)** Allocation of experimental groups. Mean ± SEM for PavCA index. Rats with similar PavCA scores were assigned to receive different treatments. Rats assigned to the orexin 1 study received either vehicle (DMSO) or the orexin 1 antagonist SB-334867 (SB). Rats assigned to the orexin 2 study received either vehicle (SAL) or the orexin 2 antagonist TCS-OX2-29 (TCS). Baseline differences in PavCA index between experimental groups were assessed by using a 2-way ANOVA with Phenotype and Treatment as independent variables and PavCA index as dependent variable. A significant effect of phenotype was found (p < 0.001). There were no other significant differences between experimental groups. N = 4-11/group for GTs, 7-16/group for STs.

For all measures analyzed, there was a significant effect of Phenotype, Session and a Phenotype x Session interaction. Relative to GTs, ST rats exhibited a greater number of lever contacts (*F*_4,118.507_ = 18.793, *p* < 0.001, Figure 3a), a greater probability to contact the lever (*F*_4,137.618_ = 29.209, *p* < 0.001, Figure 3c) and had a lower latency to approach the lever (*F*_4,155.316_ = 32.657, *p* < 0.001, Figure 3e). Post-hoc analyses revealed a significant difference between phenotypes during all 5 sessions of PavCA training (p < 0.001). During the lever-CS presentation, subjects classified as GTs showed a greater number of magazine entries (*F*_4,141.543_ = 33.338, *p* < 0.001, Figure 3b), a greater probability to enter the magazine (*F*_4,144.416_ = 39.753, *p* < 0.001, Figure 3d), and a lower latency to enter the magazine (*F*_4,138.611_ = 40.725, *p* < 0.001, Figure 3f) compared to STs. Post-hoc analyses revealed a significant difference between phenotypes for all magazine-directed measures from session 2 through session 5 (*p* < 0.05).

Since the previous measures were calculated with all subjects from Experiments 2a and 2b collapsed, it was important to ensure that the experimental groups did not differ before drug administration. To assess this, Treatment was included as a between - group variable, and acquisition data was analyzed for each experimental group. There was no significant effect of Treatment, nor were there any significant interactions with this variable, as treatment did not occur during this phase of the study. To further explore baseline differences between experimental groups, the effect of Phenotype and Treatment was assessed for the average PavCA index from sessions 4 and 5 (Figure 3h) and for lever- and magazine-directed behavior during session 5 of PavCA training, prior to treatment (Supplemental Tables 1 and 2). As expected, there was a significant effect of Phenotype (*F*_1,80_ = 965.945, *p*< 0.001), for which STs had a greater PavCA index compared to GTs rats. There was not a significant effect of Treatment, nor a significant Phenotype x Treatment interaction, suggesting that all groups were counterbalanced prior to drug testing.

### Experiment 2a-Effects of pharmacological antagonism of orexin 1 receptor on the expression of Pavlovian conditioned approach behavior and on the conditioned reinforcing properties of a Pavlovian reward-cue

Lever contacts and magazine entries are presented as the primary dependent variable in the main text, but analyses of all other lever- and magazine-directed behaviors for STs and GTs are included in Supplemental Tables 3 and 4, respectively.

#### Antagonism of OX1r in the PVT attenuates subsequent sign-tracking behavior in STs, but has no effect in GTs

To assess the effects of OX1r antagonism in the PVT on PavCA behavior, PavCA data across sessions 5, 6, and 7 were analyzed for each Phenotype. The analysis of lever contacts (i.e. sign-tracking) in STs showed no significant effect of Treatment (*F*_1,26.827_ = 2.983, *p* = 0.101) or Session (*F*_2,52.951_ = 0.568, *p* = 0.570). However, there was a significant Treatment x Session interaction (*F*_2,52.951_ = 3.735, *p* = 0.030), suggesting that the antagonism of OX1r in the PVT affected sign-tracking behavior in STs (Figure 4a). There was a trend towards a significant difference between vehicle- and drug-treated rats on session 6 (p = 0.067), when the administration of the OX1r antagonist occurred; but a more robust effect of treatment during session 7 (p = 0.023), which occurred 24 hours after drug administration. These data suggest that a single administration of the OX1r antagonist, SB-33658, may not immediately affect the attribution of incentive value to a reward-cue, but does disrupt the subsequent expression of sign-tracking behavior. There were no significant effects of Treatment (*F*_1,16.391_ = 0.318, *p* = 0.581), Session (*F*_2,16.736_ = 1.144, *p* = 0.342), nor a significant Treatment x Session interaction (*F*_2,16.736_ = 1.079, *p* = 0.363) for lever contacts in GT rats (Figure 4b), suggesting that the blockade of OX1r in the PVT has no effect on the expression of sign-tracking behavior in this phenotype.

**Figure 4.**
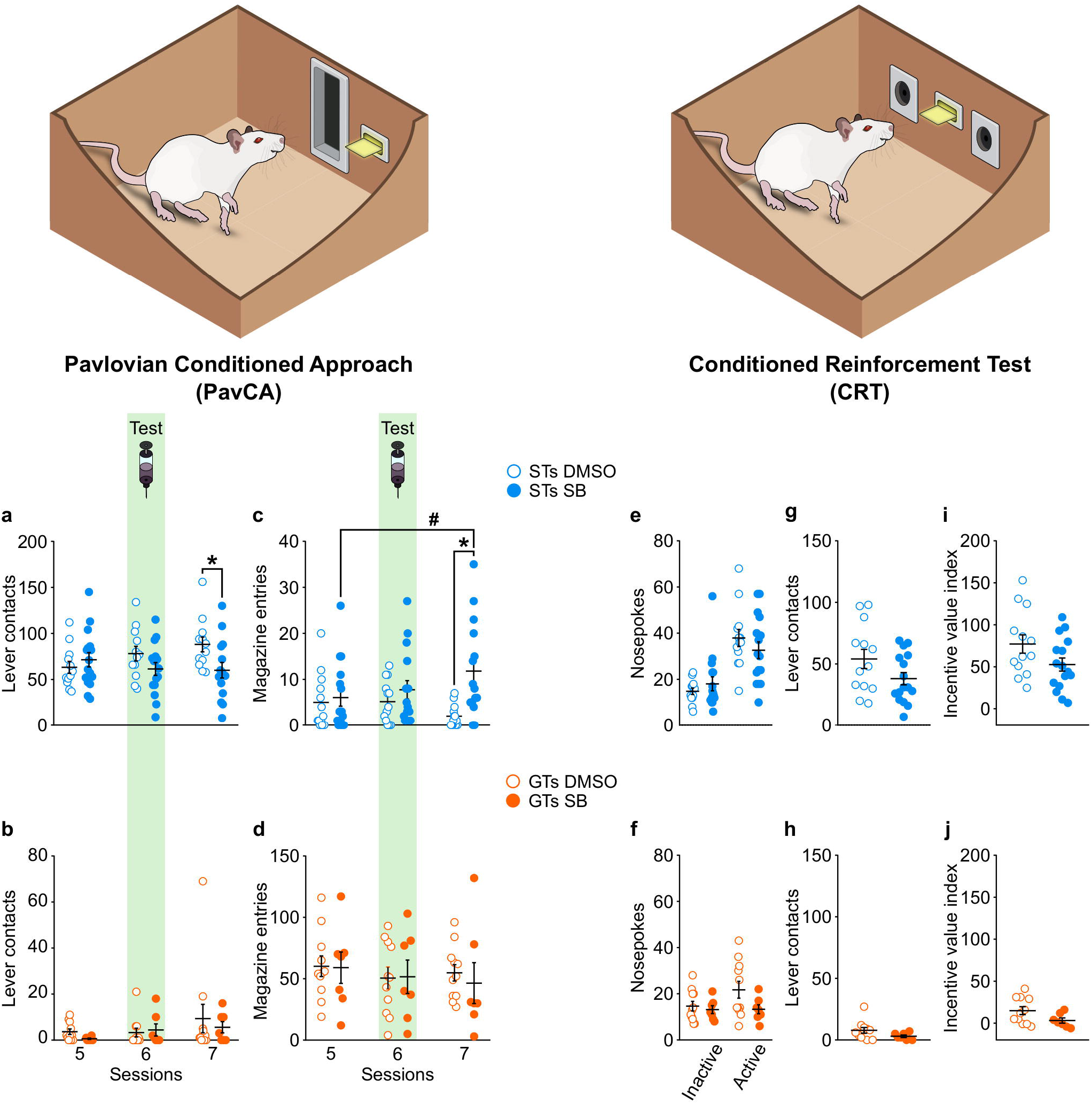
Antagonism of OX1r in the PVT prevents the escalation of sign-tracking behaviors and increases goal-tracking behavior in STs, but has no effect on the conditioning reinforcing properties of a food-cue. Left panel, mean ± SEM for **a)** number of lever contacts in sign-trackers during Session 5, 6 and 7 of PavCA training. There was a significant Treatment x Session interaction (p = 0.030). Post-hoc analyses revealed that, compared to control rats (DMSO), rats that received a single administration of the OX1r antagonist SB-334867 (SB) before Session 6 showed a decrease in lever contacts during session 7 (*p = 0.023 vs. DMSO), **c)** number of magazine entries in sign-trackers during Session 5, 6 and 7 of PavCA training. There was a significant Treatment x Session interaction (p = 0.008). Post-hoc analyses revealed that, compared to control rats (DMSO), rats that received a single administration of the OX1r antagonist SB-334867 (SB) before Session 6 showed an increase in magazine entries during session 7 (*p = 0.001 vs. DMSO; #p = 0.015 vs. Session 5). **b)** number of lever contacts and **d)** magazine entries in goal-trackers during Session 5, 6 and 7 of PavCA training. No effects were found in goal trackers. N = 13 STs-DMSO, 16 STs-SB, 11 GTs-DMSO, 7 GTs-SB. Right panel, mean ± SEM for **e)** nosepokes, **g)** lever contacts, and **i)** incentive value index in STs, and **f)** nosepokes, **h)** lever contacts, and **j)** incentive value index in GTs. There were no significant effects of OX1r antagonism in the PVT on any measures during the conditioned reinforcement test. N = 13 STs-DMSO, 16 STs-SB, 11 GTs-DMSO, 7 GTs-SB.

#### Antagonism of OX1r in the PVT increases goal-tracking behaviors in STs, but has no effect in GTs

There was not a significant effect of Treatment (*F*_1,26.909_ = 4.187, *p* = 0.051) nor Session (*F*_2,53.139_ = 0.438, *p* = 0.648) on magazine entries for STs. There was, however, a significant Treatment x Session interaction (*F*_2,53.139_ = 5.357, *p* = 0.008), suggesting that the antagonism of OX1r in the PVT affects goal-tracking behavior in STs (Figure 4c). Similar to the sign-tracking results described above, post-hoc analyses revealed a significant difference between vehicle- and drug-treated rats 24 hours after drug administration (session 7, p = 0.001), but no significant difference during session 6, immediately following the administration of the OX1r antagonist (p = 0.333). Furthermore, post-hoc analyses revealed a significant difference between session 5 and 7 in drug-treated rats (p = 0.015). Thus, despite antagonism of OX1r in the PVT having no immediate effects on goal-tracking behavior, a single administration of the OX1r antagonist SB-33658 is sufficient to promote the expression of goal-tracking behavior in STs 24 hours following drug infusion. In contrast, in GTs, there was not a significant effect of Treatment (*F*_1,16_ = 0.042, *p* = 0.840), Session (*F*_2,32_ = 1.678, *p* = 0.203), nor a significant Treatment x Session interaction (*F*_2,32_ = 0.398, *p* = 0.675) for magazine entries (Figure 4d). These findings suggest that the blockade of PVT OX1r has no effect on the expression of goal-tracking behavior in rats with an inherent tendency for this behavior.

#### Antagonism of OX-1Rs in the PVT does not alter the conditioned reinforcing properties of the lever-CS for STs or GTs

For STs, there was a significant effect of Port (F_(1,27)_ = 52.409, p = 0.000), but no effect of Treatment (F_(1,27)_ = 0.081, p = 0.778), nor a Treatment x Port interaction (F_(1,27)_ = 2.682, p = 0.113) on the number of nosepokes during the conditioned reinforcement test. An unpaired t-test showed a trend towards a significant effect of Treatment for the number of lever contacts (t_(27)_ = 1.802, p = 0.083; Figure 6c) and incentive value index (t_(27)_ = 1.876, p = 0.071; Figure 6e) for STs. For GTs, there was not a significant effect of Port (F_(1,16)_ = 1.833, p = 0.195), Treatment (F_(1,16)_ = 2.706, p = 0.119), nor a Treatment x Port interaction (F_(1,16)_ = 1.691, p = 0.212; Figure 6b). In addition, GTs showed no effect of Treatment for lever contacts (t_(16)_ = 1.614, p = 0.126) and there was a trend towards significance for the incentive value index (t_(16)_ = 1.818, p = 0.088; Figures 6d,f). These results suggest that OX1r antagonism in the PVT does not significantly alter the conditioned reinforcing properties of the lever-CS for either STs or GTs.

### Experiment 2b-Effects of pharmacological antagonism of OX2r on the expression of Pavlovian conditioned approach behavior and the conditioned reinforcing properties of a Pavlovian reward-cue

Lever contacts and magazine entries are presented as the primary dependent variable in the main text, but analyses of all other lever- and magazine-directed behaviors for STs and GTs are included in Supplemental Tables 5 and 6, respectively.

#### Antagonism of OX2r in the PVT prevents the escalation of sign-tracking behaviors in STs, but has no effect in GTs

There was a significant effect of Session (*F*_2,24.730_ = 4.261, *p* = 0.026) and a significant Treatment x Session interaction (*F*_2,24.730_ = 6.937, *p* = 0.004) for the number of lever contacts in STs, suggesting that antagonism of OX2r in the PVT affects sign-tracking behavior. Post-hoc analyses revealed a significant difference between vehicle- and drug-treated rats on session 6 (p = 0.039), during which antagonism of OX2r in the PVT reduced sign-tracking behavior compared to vehicle-treated controls of the same phenotype (Figure 5a). In addition, post-hoc comparisons revealed a significant difference between session 5, 6 and 7 in vehicle-treated rats, but not in drug-treated rats. These data suggest that a single administration of the OX2r antagonist, TCS-OX2-29, is sufficient to prevent the increase in sign-tracking behavior shown by control rats (Figure 5a). While there was a significant effect of Session (*F*_2,9.740_ = 4.837, *p* = 0.035; Figure 5b) in GTs, there was not a significant effect of Treatment (*F*_1,9.016_ = 0.704, *p* = 0.423), nor a significant Treatment x Session interaction (*F*_2,9.740_ = 0.370, *p* = 0.700). Together, these results indicate that pharmacological antagonism of OX2r in the PVT prevents the escalation of sign-tracking behavior in STs, but has no effect in GTs.

**Figure 5.**
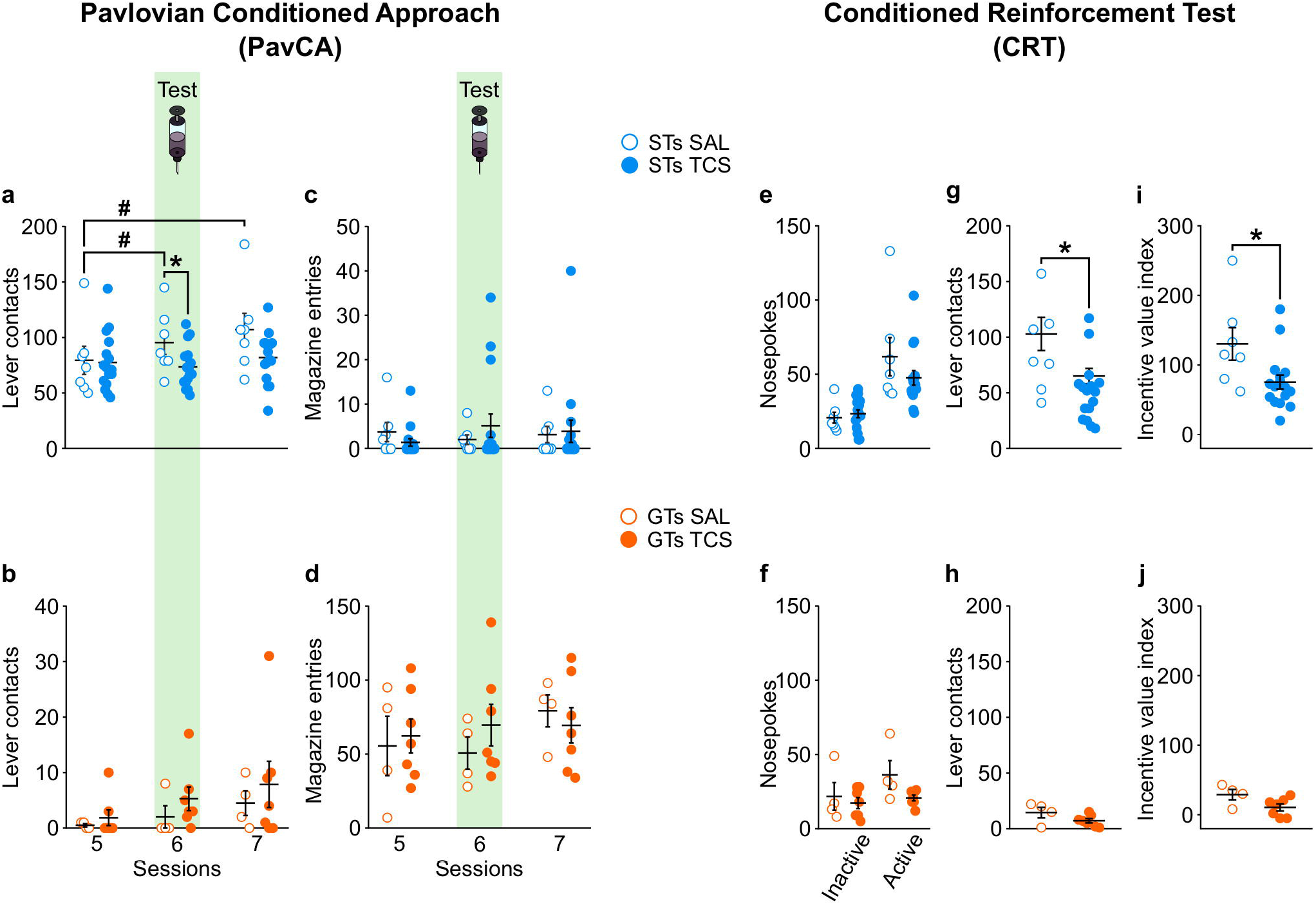
Antagonism of OX2r in the PVT prevents the escalation of sign-tracking behaviors and reduces the conditioned reinforcing properties of a reward-paired cue in sign-trackers, but has no effect in GTs. Left panel, mean ± SEM for **a**) number of lever contacts in sign-trackers during Session 5, 6 and 7 of PavCA training. There was a significant Treatment x Session interaction (p = 0.004). Post-hoc analyses revealed that, compared to control rats (SAL), rats that received a single administration of the OX2r antagonist TCS-OX2-29 (TCS) before Session 6 showed a decrease in lever contacts (*p = 0.039 vs. SAL). **c)** Number of magazine entries in sign-trackers during Session 5, 6 and 7 of PavCA training. No effects were found for magazine entries in sign-trackers. **b)** number of lever contacts and **d)** magazine entries in goal-trackers during Session 5, 6 and 7 of PavCA training. There were no significant effects in goal trackers. N = 7 STs-SAL, 16 STs-TCS, 4 GTs-SAL, 7 GTs-TCS. Right panel, mean ± SEM for **e)** nosepokes, **g)** lever contacts, and **i)** incentive value index in STs. A significant reduction in lever contacts and incentive index was apparent in STs who received infusion of the OX2r antagonist TCS OX2 29 into the PVT (vs. controls). For GTs, there was no significant effect of drug administration for **f)** nosepokes, **h)** lever contacts, or the **j)** incentive value index. N = 7 STs-SAL, 16 STs-TCS, 4 GTs-SAL, 7 GTs-TCS.

#### Antagonism of OX2r in the PVT does not affect goal-tracking behaviors

There was not a significant effect of OX2r antagonism in the PVT on magazine entries in STs (Figure 5c), as there was no effect of Session (*F*_2,21_ = 0.283, *p* = 0.757), Treatment (*F*_1,21_ = 0.027, *p* = 0.871), nor a significant Treatment x Session interaction (*F*_2,21_ = 1.447, *p* = 0.258). Similarly, in GTs, there were no significant effects of Session (*F*_2,17.943_ = 2.803, *p* = 0.087) or Treatment (*F*_1,9.392_ = 0.084, *p* = 0.778), and just a trend towards a significant Treatment x Session interaction (*F*_2,17.943_ = 3.062, *p* = 0.072; Figure 5d). Thus, antagonism of OX2r in the PVT does not significantly affect goal-tracking behavior in either STs or GTs.

#### Antagonism of OX-2Rs in the PVT affects the conditioned reinforcing properties of the lever-CS in STs

When assessing the effects of OX2r antagonism on nosepokes in the CRT paradigm, there was a significant effect of Port (*F*_1,21_ = 43.666, *p* = 0.000), but no effect of Treatment (F_(1,21)_ = 0.695, p = 0.414), nor a Treatment x Port interaction (F_(1,21)_ = 2.941, p = 0.101, Figure 5e) for STs. There was, however, a significant effect of OX2r antagonism on the number of lever contacts in STs (t_(21)_ = 2.664, p = 0.015; Figure 5g). Even though pokes into the active port result in only a brief 2-s presentation of the lever-CS, vehicle-treated rats tended to engage with the lever-CS to a greater extent than drug-treated rats during the CRT session. In agreement, compared to vehicle-treated rats, antagonism of OX2r in the PVT reduced the incentive value index in STs (t_(21)_ = 2.546, p = 0.019; Figure 5i). For GTs, there was a significant effect of Port (*F*_1,9_ = 11.078, *p* = 0.009), but no effect of Treatment (*F*_1,9_ = 1.817, *p* = 0.211), and only a trend towards significance for a Treatment x Port interaction (*F*_1,9_ = 4.225, *p* = 0.070; Figure 5f). There was not a significant effect of Treatment for lever contacts (t_(9)_ = 1.710, p = 0.121) or the incentive value index (t_(9)_ = 2.047, p = 0.089) in GTs during CRT (Figure 5h,j).

## Discussion

We performed two separate experiments to examine both the role of the LHA and orexin signaling within the PVT in the attribution of incentive salience to a Pavlovian-conditioned food cue. In Experiment 1, we tested a causal influence of the LHA on the acquisition of sign- and goal-tracking behavior. A bilateral excitotoxic lesion of the LHA before Pavlovian training resulted in an attenuation of lever-directed behaviors during PavCA training, without affecting magazine-directed behaviors (Figure 2). These results indicate that an intact LHA is required for the acquisition of sign-tracking, but not goal-tracking behavior. Originally considered to be the feeding center of the brain (Anand & Brobeck, 1951), the LHA has long been recognized to play a role in motivated and reward-related behaviors (Margules & Olds, 1962; Devenport & Balagura, 1971; DiLeone et al., 2003; Nieh et al., 2016; Stuber & Wise, 2016; Tyree & de Lecea, 2017). While the LHA has previously been shown to respond to or be activated by reward-paired and incentive-motivational cues (Nieh et al., 2015; Haight et al., 2017), this is the first study to causally link the LHA to the attribution of incentive salience to reward cues. Moreover, the fact that goal-tracking behavior was not affected, suggests that the LHA is not critical for the attribution of predictive value to reward cues. These findings are in agreement with our prior research, demonstrating that, relative to GTs, STs have greater c-fos counts in the LHA, and predominantly in neurons of the LHA that project to the PVT (Haight et al., 2017). Thus, the LHA appears to play a critical role in incentive motivational processes, and likely does so via the PVT.

The findings described above were expanded in Experiment 2, where we examined the role of orexinergic activity in the PVT on the attribution of incentive salience to a food-cue. To this end, we tested the effects of selective antagonism of either the OX1r (Experiment 2a) or OX2r (Experiment 2b) in the PVT on Pavlovian conditioned approach behavior and the conditioned reinforcing properties of the food-cue. Unlike Experiment 1, this study focused on the expression of Pavlovian conditioned approach behavior, after the conditioned responses had been acquired. We found that the OX2r antagonist immediately decreased the incentive-motivational value of the reward-cue when administered directly into the PVT of ST rats, evidenced by decreased lever contacts during the PavCA paradigm. Thus, orexin signaling at OX2r receptors in the PVT appears to be directly involved in mediating approach behavior directed towards an incentive-motivational stimulus. By blocking this signal, the incentive-motivational value of the cue is reduced, and a deficit in approach behavior can be readily observed. In addition, performance decrements in PavCA behavior were only observed in rats that developed a sign-tracking phenotype, and were not apparent in rats that developed a goal-tracking phenotype. Presumably, this is because GTs do not assign incentive motivational value to the reward cue (Robinson & Flagel, 2009), further indicating that the decrease in behavior observed in STs was due to a loss in the incentive motivational value of the lever-CS, and not a general disruption of the stimulus-reward association.

Antagonism of OX1r receptors in the PVT did not immediately attenuate PavCA behavior. Instead, the observable effects of a single intra-PVT injection of the OX1r receptor antagonist were largely apparent 24 hours after drug administration (session 7). On session 7, there was both a decrease in lever-directed behavior (sign-tracking), and an increase in food-cup directed behavior (goal-tracking) in ST rats. Thus, blockade of OX1 receptors in the PVT appeared to disrupt the sign-tracking trajectory and permit STs to adapt a predictive learning strategy on subsequent sessions. In contrast, blockade of OX2r receptors in the PVT immediately inhibited lever-directed behaviors in STs, without affecting magazine-directed behaviors. Orexin, therefore, appears to have multiple roles in the PVT, with different receptor subtypes mediating distinct aspects of stimulus-reward processing. Speculatively, activity at OX2r may integrate incentive-salience signals from the LHA to the PVT in real time, while activity at OX1r may encode long-term incentive values associated with Pavlovian-conditioned cues. Thus, disruption of OX1r signaling does not manifest itself in behavior immediately, but only becomes apparent when the stimulus is presented the following day. This could also potentially explain why there was a concomitant increase in goal-tracking behavior. The cue-reward association remains intact, and it still able to elicit a conditioned response, but the long-term incentive value of the cue has been reduced, and the focus of the response shifts towards the location of reward delivery. Additional studies will be needed to further explore these diverging roles for orexin signaling.

In addition to PavCA behavior, the effects of orexin antagonism on behavior during a conditioned reinforcement test were assessed. The conditioned reinforcement test was performed as a second measure of the incentive value of the reward cue, since previous work has demonstrated that it can be quite difficult to interrupt an already-acquired sign-tracking phenotype, which is resistant to extinction (Ahrens et al., 2016; Fitzpatrick et al., 2019). There was no effect of administration of the OX1r antagonist, on the conditioned reinforcing properties of the food cue. Similarly, administration of the OX2r antagonist did not affect instrumental responding for the food cue; but it did affect approach to the cue when it was presented and thereby the incentive value index. These findings are in agreement with those described above, as blockade of OX2r in the PVT appears to attenuate cue-directed approach behaviors during either Pavlovian training or a conditioned reinforcement test. Thus, OX2r signaling in the PVT appears to encode the incentive motivational value of reward cues, but may not be directly involved in the conditioned reinforcing properties. These findings are not necessarily discordant with one another, as different neural mechanisms underlie Pavlovian vs. instrumental responding for reward cues (Cardinal et al., 2002; Yin et al., 2008).

The data reported here add to a growing literature highlighting an important role for the LHA-PVT circuit in motivated behaviors and cue-reward learning (Kirouac et al., 2005; Martin-Fardon & Boutrel, 2012; James & Dayas, 2013; Matzeu et al., 2014; Haight et al., 2017). Unlike previous studies, however, we were able to isolate the role of the LHA and orexin signaling in the PVT specifically in incentive motivational processes. Recently, Otis and colleagues demonstrated that projection cells from the PFC to the PVT send signals encoding cue-reward relationships, while projection cells from the LHA to the PVT (putatively GABAergic) send consummatory signals to the PVT (Otis et al., 2019). While 100% of the PVT neurons observed by Otis had an inhibitory response following LHA stimulation, 35% of them also showed excitatory responses. Orexin is an excitatory peptide that has been shown to increase activity of PVT neurons (Kolaj et al., 2014). Speculatively, a proportion of the LHA inputs targeted by Otis and colleagues were likely orexinergic, and, according to our data, send signals to the PVT encoding the incentive-motivational value of the reward-paired CS. The LHA-PVT circuit, therefore, is likely not limited to consummatory signals. Rather, we speculate that, in the study described by Otis et al. (2019), the excitatory orexinergic activity that encodes the incentive motivational value of a reward cue is interwoven within the larger GABAergic input from the LHA to the PVT. Further, in agreement with Otis and colleagues, we believe that output from the PVT to the nucleus accumbens is a critical component of reward processing. We propose, however, that this circuit is especially important for incentive motivational processes. In support, our prior findings show that neurons projecting from the PVT to the NAc are activated to a greater extent in sign-trackers than goal-trackers in response to a Pavlovian food cue (Haight et al., 2017). That is, these neurons are engaged by an incentive stimulus, but not a predictive stimulus. In addition, we have long-known that stimulation of PVT-NAc projections can elicit nucleus accumbens dopamine release, and can do so independent of the ventral tegmental area (Parsons et al., 2007). As we know that dopamine release in the NAc is critical for sign-tracking, but not goal-tracking, behavior, we propose that the PVT-NAc pathway is the final integrative component of a hypothalamic (LHA) – thalamic (PVT) – striatal circuit mediating the incentive value of reward cues. To determine if this is indeed the case, future studies will need to combine elegant technical approaches, like that used by Otis and colleagues, with behavioral outcome measures that permit the dissociation of complex psychological phenomenon, like those used here.

There are two potential limitations of the studies conducted here. First is the fact that a food stimulus was used, and both the LHA and the PVT have been shown to be components of the ingestive circuitry (Stratford & Wirtshafter, 2013; Cheng et al., 2018). In terms or orexinergic involvement, previous work has demonstrated that administration of orexin-A into the posterior PVT leads to increased consumption of 2% sucrose solution (Barson et al., 2015), while knockdown of OX-1Rs in the PVT reduces hedonic feeding of high-fat chow in rats (Choi et al., 2012). While the impact of this circuit on food consumption is important, we do not think that it affected the current results. The fact that subjects continued to retrieve and consume food pellets during PavCA testing suggests that the desire to consume the food reward was not affected. Second, orexinergic inputs to the PVT have been recently identified as part of the circuity promoting wakefulness via signals to the NAc (Ren et al., 2018), demonstrating yet another role for this diverse and heterogeneous circuit. However, we do not believe this confounded the reported results. First, Ren and colleagues found that PVT manipulations had minimal effects on reducing wakefulness during the light cycle, when our animals were tested. In addition, if blockade of OX1r or OX2r in the PVT reduced wakefulness, then we would have expected to observe a decrease in PavCA behavior in both STs and GTs. Instead, we only observed selective behavioral deficits that were tied to incentive salience attribution. Thus, we believe we have identified an additional and novel role for orexinergic signaling in the PVT.

In conclusion, this work supports the notion that the attribution of incentive salience to reward cues is mediated, predominantly, by bottom-up pathways. Specifically, we identify a novel role for orexinergic signaling in the PVT that appears to be originating from the lateral hypothalamic area. In addition, we have recognized potentially distinct roles for orexin activity at OX1r and OX2r within the PVT, but believe their convergent properties act to encode the incentive motivational value of reward cues. While further work is needed to fully understand the complex role of orexin transmission in the PVT and the ability of the LHA to evoke these downstream effects, this study identifies a novel role for the LHA and orexin signaling within the PVT in incentive motivational processes.

## Acknowledgements

This work was supported by the National Institute on Drug Abuse branch of the National Institutes of Health (T32DA007268 [IRC], F31DA037680 [JLH], T32DA007821 [BNK], K08DA037912 & R01DA044960 [JDM], R01DA038599 [SBF]) and The National Science Foundation (GRFP [CMR]).

---

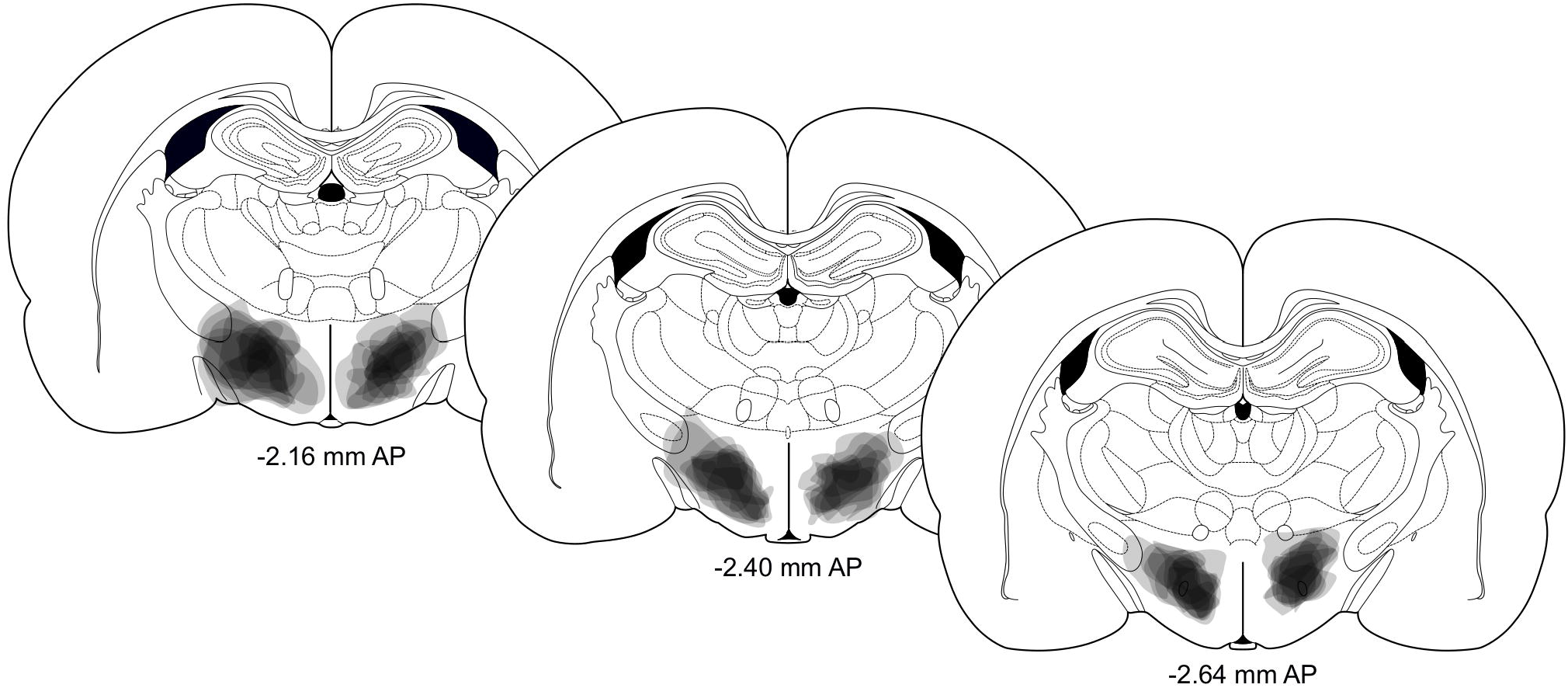

