## Supplemental tables 1, 2, 3, 4, 5, 6 for "The lateral hypothalamus and orexinergic transmission in the paraventricular thalamus promote the attribution of incentive salience to reward-associated cues"

**Supplemental data**

**Supplemental Table 1.** The effect of phenotype and treatment on lever- and magazine-directed behaviors during session 5 of PavCA training for Experiment 2a.

|  | Experiment 2a: Antagonism of orexin 1 receptors in the PVT | | | | | | | | |
| --- | --- | --- | --- | --- | --- | --- | --- | --- | --- |
|  | **Lever-directed behaviors** | | | | | | | | |
|  | Lever contacts | | | Probability lever | | | Latency lever | | |
|  | DF | F | p | DF | F | p | DF | F | p |
| Phenotype | 1,43 | 55.907 | **0.000** | 1,43 | 383.553 | **0.000** | 1,43 | 121.863 | **0.000** |
| Treatment | 1,43 | 0.121 | 0.729 | 1,43 | 0.281 | 0.599 | 1,43 | 0.020 | 0.888 |
| Phenotype*Treatment | 1,43 | 0.001 | 0.979 | 1,43 | 1.864 | 0.179 | 1,43 | 1.153 | 0.289 |
|  | **Magazine-directed behaviors** | | | | | | | | |
|  | Magazine entries | | | Probability magazine | | | Latency magazine | | |
|  | DF | F | p | DF | F | p | DF | F | p |
| Phenotype | 1,43 | 84.412 | **0.000** | 1,43 | 79.642 | **0.000** | 1,43 | 72.540 | **0.000** |
| Treatment | 1,43 | 0.000 | 0.992 | 1,43 | 0.211 | 0.648 | 1,43 | 0.599 | 0.443 |
| Phenotype*Treatment | 1,43 | 0.036 | 0.850 | 1,43 | 1.341 | 0.253 | 1,43 | 0.812 | 0.373 |

**Supplemental Table 2.** The effect of phenotype and treatment on lever- and magazine-directed behaviors during session 5 of PavCA training for Experiment 2b.

|  | Experiment 2b: Antagonism of orexin 2 receptors in the PVT | | | | | | | | |
| --- | --- | --- | --- | --- | --- | --- | --- | --- | --- |
|  | **Lever-directed behaviors** | | | | | | | | |
|  | Lever contacts | | | Probability lever | | | Latency lever | | |
|  | DF | F | p | DF | F | p | DF | F | p |
| Phenotype | 1,30 | 71.177 | **0.000** | 1,30 | 406.365 | **0.000** | 1,30 | 128.276 | **0.000** |
| Treatment | 1,30 | 0.001 | 0.978 | 1,30 | 0.076 | 0.785 | 1,30 | 0.000 | 0.992 |
| Phenotype*Treatment | 1,30 | 0.031 | 0.862 | 1,30 | 0.210 | 0.650 | 1,30 | 0.033 | 0.857 |
|  | **Magazine-directed behaviors** | | | | | | | | |
|  | Magazine entries | | | Probability magazine | | | Latency magazine | | |
|  | DF | F | p | DF | F | p | DF | F | p |
| Phenotype | 1,30 | 59.472 | **0.000** | 1,30 | 87.647 | **0.000** | 1,30 | 62.383 | **0.000** |
| Treatment | 1,30 | 0.093 | 0.763 | 1,30 | 0.003 | 0.958 | 1,30 | 0.074 | 0.788 |
| Phenotype*Treatment | 1,30 | 0.390 | 0.537 | 1,30 | 0.785 | 0.383 | 1,30 | 1.260 | 0.271 |

**Supplemental Table 3.** The effect of treatment and session on lever- and magazine-directed behaviors across sessions 5, 6 and 7 of PavCA for sign-tracker rats in Experiment 2a.

|  | Experiment 2a STs: Antagonism of orexin 1 receptors in the PVT | | | | | | | | |
| --- | --- | --- | --- | --- | --- | --- | --- | --- | --- |
|  | **Lever-directed behaviors** | | | | | | | | |
|  | Lever contacts | | | Probability lever | | | Latency lever | | |
|  | DF | F | p | DF | F | p | DF | F | p |
| Treatment | 1,26.827 | 2.893 | 0.101 | 1,26.719 | 2.090 | 0.160 | 1,27.763 | 0.302 | 0.587 |
| Session | 2,52.951 | 0.568 | 0.570 | 2,53.039 | 0.095 | 0.910 | 2,52.215 | 0.477 | 0.623 |
| Treatment*Session | 2,52.951 | 3.735 | **0.030** | 2,53.039 | 2.836 | 0.068 | 2,52.215 | 2.610 | 0.083 |
|  | **Magazine-directed behaviors** | | | | | | | | |
|  | Magazine entries | | | Probability magazine | | | Latency magazine | | |
|  | DF | F | p | DF | F | p | DF | F | p |
| Treatment | 1,26.909 | 4.187 | 0.051 | 1,26.865 | 6.006 | **0.021** | 1,26.936 | 4.094 | 0.053 |
| Session | 2,53.139 | 0.438 | 0.648 | 2,53.037 | 0.812 | 0.450 | 2,53.173 | 1.112 | 0.337 |
| Treatment*Session | 2,53.139 | 5.357 | **0.008** | 2,53.037 | 5.839 | **0.005** | 2,53.173 | 4.888 | **0.011** |

**Supplemental Table 4.** The effect of treatment and session on lever- and magazine-directed behaviors across sessions 5, 6 and 7 of PavCA for goal-tracker rats in Experiment 2a.

|  | Experiment 2a GTs: Antagonism of orexin 1 receptors in the PVT | | | | | | | | |
| --- | --- | --- | --- | --- | --- | --- | --- | --- | --- |
|  | **Lever-directed behaviors** | | | | | | | | |
|  | Lever contacts | | | Probability lever | | | Latency lever | | |
|  | DF | F | p | DF | F | p | DF | F | p |
| Treatment | 1,16.391 | 0.318 | 0.581 | 1,16.259 | 0.070 | 0.795 | 1,16.838 | 0.644 | 0.434 |
| Session | 2,16.736 | 1.144 | 0.342 | 2,20.961 | 2.657 | 0.094 | 2,19.533 | 0.982 | 0.392 |
| Treatment*Session | 2,16.736 | 1.079 | 0.363 | 2,20.961 | 1.685 | 0.210 | 2,19.533 | 1.070 | 0.362 |
|  | **Magazine-directed behaviors** | | | | | | | | |
|  | Magazine entries | | | Probability magazine | | | Latency magazine | | |
|  | DF | F | p | DF | F | p | DF | F | p |
| Treatment | 1,16 | 0.042 | 0.840 | 1,16 | 1.418 | 0.251 | 1,16 | 0.836 | 0.374 |
| Session | 2,32 | 1.678 | 0.203 | 2,32 | 0.047 | 0.954 | 2,32 | 0.308 | 0.737 |
| Treatment*Session | 2,32 | 0.398 | 0.675 | 2,32 | 0.973 | 0.389 | 2,32 | 0.561 | 0.576 |

**Supplemental Table 5.** The effect of treatment and session on lever- and magazine-directed behaviors across sessions 5, 6 and 7 of PavCA for sign-tracker rats in Experiment 2b.

|  | Experiment 2b STs: Antagonism of orexin 1 receptors in the PVT | | | | | | | | |
| --- | --- | --- | --- | --- | --- | --- | --- | --- | --- |
|  | **Lever-directed behaviors** | | | | | | | | |
|  | Lever contacts | | | Probability lever | | | Latency lever | | |
|  | DF | F | p | DF | F | p | DF | F | p |
| Treatment | 1,21.101 | 2.377 | 0.138 | 1,21 | 0.534 | 0.473 | 1,21 | 1.045 | 0.318 |
| Session | 2,24.730 | 4.261 | **0.026** | 2,42 | 0.968 | 0.388 | 2,42 | 5.074 | **0.016** |
| Treatment*Session | 2,24.730 | 6.937 | **0.004** | 2,42 | 1.480 | 0.239 | 2,42 | 3.520 | **0.048** |
|  | **Magazine-directed behaviors** | | | | | | | | |
|  | Magazine entries | | | Probability magazine | | | Latency magazine | | |
|  | DF | F | p | DF | F | p | DF | F | p |
| Treatment | 1,21 | 0.027 | 0.871 | 1,21 | 0.017 | 0.899 | 1,21.098 | 0.000 | 0.994 |
| Session | 2,21 | 0.283 | 0.757 | 2,21 | 0.022 | 0.978 | 2,31.892 | 0.114 | 0.892 |
| Treatment*Session | 2,21 | 1.447 | 0.258 | 2,21 | 2.300 | 0.125 | 2,31.892 | 2.999 | 0.064 |

**Supplemental Table 6.** The effect of treatment and session on lever- and magazine-directed behaviors across sessions 5, 6 and 7 of PavCA for goal-tracker rats in Experiment 2b.

|  | Experiment 2b GTs: Antagonism of orexin 1 receptors in the PVT | | | | | | | | |
| --- | --- | --- | --- | --- | --- | --- | --- | --- | --- |
|  | **Lever-directed behaviors** | | | | | | | | |
|  | Lever contacts | | | Probability lever | | | Latency lever | | |
|  | DF | F | p | DF | F | p | DF | F | p |
| Treatment | 1,9.016 | 0.704 | 0.423 | 1,9.022 | 0.936 | 0.358 | 1,9.384 | 0.279 | 0.610 |
| Session | 2,9.740 | 4.837 | **0.035** | 2,11.890 | 5.737 | **0.018** | 2,18 | 3.166 | 0.066 |
| Treatment*Session | 2,9.740 | 0.370 | 0.700 | 2,11.890 | 0.583 | 0.574 | 2,18 | 0.086 | 0.918 |
|  | **Magazine-directed behaviors** | | | | | | | | |
|  | Magazine entries | | | Probability magazine | | | Latency magazine | | |
|  | DF | F | p | DF | F | p | DF | F | p |
| Treatment | 1,9.391 | 0.084 | 0.778 | 1,9.036 | 0.042 | 0.842 | 1,9.283 | 0.427 | 0.529 |
| Session | 2,17.943 | 2.803 | 0.087 | 2,12.363 | 2.037 | 0.172 | 2,17.844 | 0.348 | 0.711 |
| Treatment*Session | 2,17.943 | 3.062 | 0.072 | 2,12.363 | 3.707 | 0.055 | 2,17.844 | 1.187 | 0.328 |

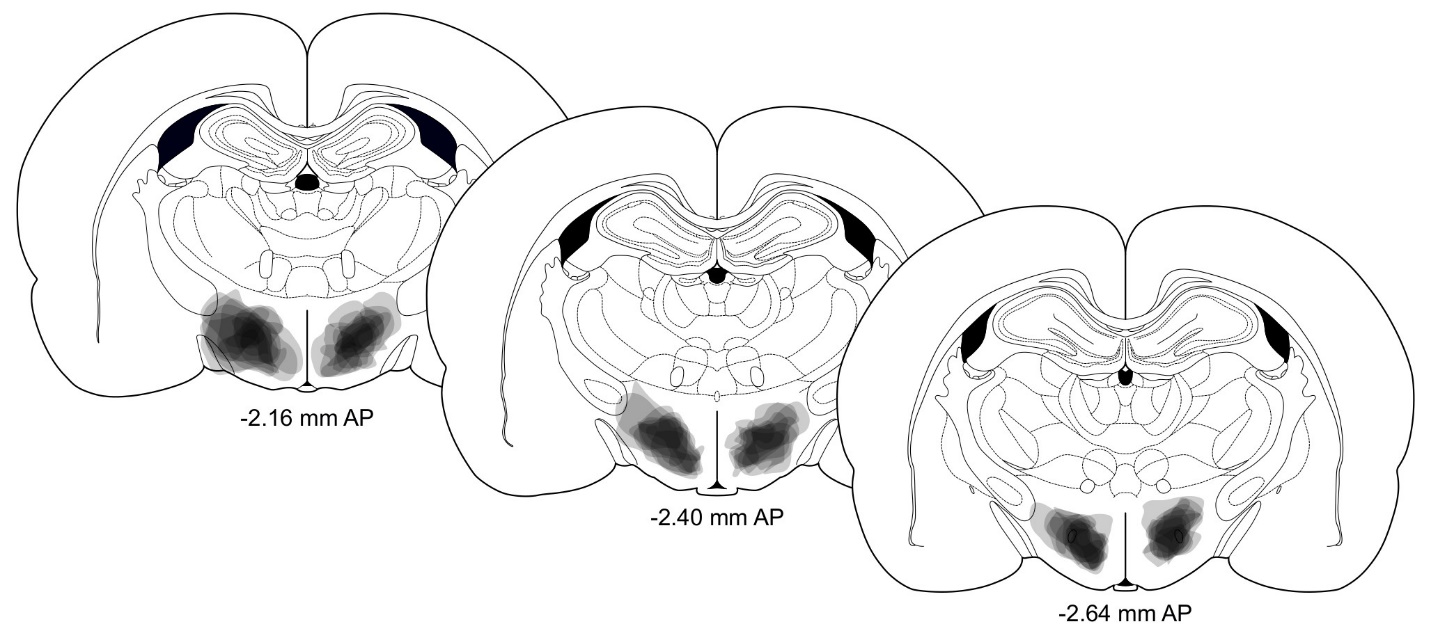

**Supplemental Figure 1.** Representative schematics of the spread of NMDA infusion in the LHA-lesioned rats.
